# Identifying Drug Sensitivity Subnetworks with NETPHIX

**DOI:** 10.1101/543876

**Authors:** Yoo-Ah Kim, Rebecca Sarto Basso, Damian Wojtowicz, Amanda Liu, Dorit S. Hochbaum, Fabio Vandin, Teresa M. Przytycka

## Abstract

Phenotypic heterogeneity in cancer is often caused by different patterns of genetic alterations. Understanding such phenotype-genotype relationships is fundamental for the advance of personalized medicine. One of the important challenges in the area is to predict drug response on a personalized level and to understand the causes of different responses. The pathway-centric view of cancer significantly advanced the understanding of genotype-phenotype relationships. However, most network identification methods in cancer focus on identifying subnetworks that include general cancer drivers or are associated with discrete features such as cancer subtypes, hence cannot be applied directly for the analysis of continuous features like drug response. On the other hand, existing genome wide association approaches do not fully utilize the complex and heterogeneous proprieties of cancer mutational landscape. To address these challenges, we developed a computational method, named NETPHIX (NETwork-to-PHenotype assocIation with eXclusivity), which aims to identify subnetworks of genes whose genetic alterations are associated with a continuous cancer phenotype. Leveraging the properties of cancer mutations such as mutual exclusivity and the interactions among genes, we formulate the problem as an integer linear program and solve it optimally to obtain a set of associated genes. Applied to a large-scale drug screening dataset, NETPHIX uncovered gene modules significantly associated with drug responses, and many of the modules are also validated in another independent dataset. Utilizing interaction information, NETPHIX modules are functionally coherent, and can thus provide important insights into drug action.

## 1 Introduction

Genetic alterations in cancer are associated with diverse phenotypic properties such as drug response or patient survival. However, the identification of mutations causing specific phenotypes and the interpretation of the phenotype-genotype relationships remain challenging due to a large number of passenger mutations and cancer heterogeneity. Indeed, the relationships between genotype and phenotype in most tumors are complex and different mutations in functionally related genes can lead to the same phenotype. The pathway-centric view of cancer [1, 2, 3] suggests that cancer phenotype should be considered from the context of dysregulated pathways rather than from the perspective of mutations in individual genes. Such a pathway-centric view significantly advanced the understanding of the mechanisms of tumorigenesis. Many computational methods to identify cancer driving mutations have been developed based on pathway-centric approaches [4, 5, 6, 7, 8, 9]. Network based approaches have been further applied to find subnetworks associated with various disease phenotypes [4, 6, 10, 11, 12]. Those methods have been developed aiming to find genes whose mutations are associated specifically with given phenotypes rather than finding general cancer drivers.

Recent projects have characterized drug sensitivity in hundreds of cancer cell lines for a large number of drugs [13, 14]. This data, together with information about the genetic alterations in these cells, can be used to understand how genomic alterations impact drug sensitivity. While the success of network based methods in other cancer domains suggests that such approaches should be also useful in the studies of drug response, most of previous approaches focused on discrete phenotypic traits – e.g., cancer vs. healthy, good or bad prognosis, or cancer subtypes – and therefore, cannot be directly applied to the analysis of continuous features such as drug sensitivity.

To address these challenges, we introduce a computational tool named NETPHIX (NETwork-to-PHenotype assocIation with eXclusivity). With the goal of identifying mutated subnetworks that are associated with a continuous phenotype, our algorithm builds on combinatorial optimization techniques involving *connected set cover*. The objective function of NETPHIX allows to find subnetworks with a mix of genes associated with increased or increased sensitivity simultaneously, considering interactions between resistance and sensitivity alterations. In addition, we designed the objective function to preferentially select mutually exclusive genes in the solution, utilizing an observation that cancer related mutations tend to be mutually exclusive [15, 16, 17, 7, 18, 19]. We formulate the problem as an integer linear program and solve it to optimality using CPLEX. This approach together with selecting significantly associated modules allows to leave out passenger mutations from the sensitivity networks.

Several algorithms have been previously developed for the identification of mutations associated with drug response [20, 21, 22] but without considering functional relationships among genes. For example, REVEALER used a re-scaled mutual information metric to iteratively identify a set of genes associated with the phenotype [21]. UNCOVER employs an integer linear programming formulation based on the set cover problem, by designing the objective function to maximize the association with the phenotype and preferentially select mutually exclusive gene sets [20]. While UNCOVER uses a similar objective function as NETPHIX, it does not allow to pick up mixed sensitivity modules nor utilize network information. LOBICO [22, 23] is designed to identify a set of genes whose alterations are associated with differences in drug response. The algorithm is formulated as an integer linear program, based on logic models of binary input features that explain a continuous phenotype variable. However, none of the algorithms mentioned above utilize network interaction information. Since perturbations in functionally related genes are likely to lead to similar phenotypes, functional interaction information can be helpful for the identification of phenotype associated genes.

There have been related studies combining GWAS analysis with network constraints [24, 25, 26, 27]. While these methods generally perform well at broadly pointing to disease related genes, they do not consider complex properties of cancer mutations such as the aforementioned mutual exclusivity of cancer drivers, and are not designed to zoom in on subnetworks that are specific enough to help understand drug action. As discussed later, the genomic landscape related to drug response can be complex and mutations in different genes in the same pathway can affect the response differently. Pharmaceutical drugs are often developed to target specific genes, and the response depends on the function and the mutation status of the gene as well as other genes in the same pathway.

We evaluated NETPHIX and other related methods using simulations and showed that NETPHIX outperforms all the competing methods. Applying NETPHIX to a large scale drug response data (Genomics of Drug Sensitivity in Cancer(GDSC)), we identified sensitivity-associated subnetworks for many of the drugs. We were also able to validate many of the identified modules with an indenpendent drug screening datase (The Cancer Therapeutics Response Portal (CTRP)). These subnetworks provided important insights into drug action. Effective computational methods to discover these associations will improve our understanding of the molecular mechanism of drug sensitivity, help to identify potential dug combinations, and have a profound impact on genome-driven, personalized drug therapy. NETPHIX is available at https://www.ncbi.nlm.nih.gov/CBBresearch/Przytycka/index.cgi#netphix

## 2 Results

### 2.1 NETPHIX overview

Given gene alteration information and drug sensitivity profiles (or any cancer-related, continuous phenotype) for the same set of cancer samples (or cell lines), NETPHIX aims to identify genetic alterations underlying the phenotype of interest (Fig. 1a). Based on the assumption that genes whose mutations lead to the same phenotype must be functionally related, NETPHIX also utilizes functional interaction information among genes as an input, and enforces the identified genes to be highly *connected* in the network. The problem is formulated as an integer linear program (ILP) based on a connected set cover approach. Below we briefly describe the connected set cover based algorithm. For the formal definition of the problem and the detailed ILP formulation, see Section S1.1 and S1.2, respectively. By running ILP instances with different parameters and obtaining optimal solutions using CPLEX [28], we generate candidate modules whose aggregate alterations may be associated with a given drug response. Statistical significance of candidate modules is then assessed with a permutation test, and a set of maximal modules are selected as final sensitivity modules.

**Figure 1:**
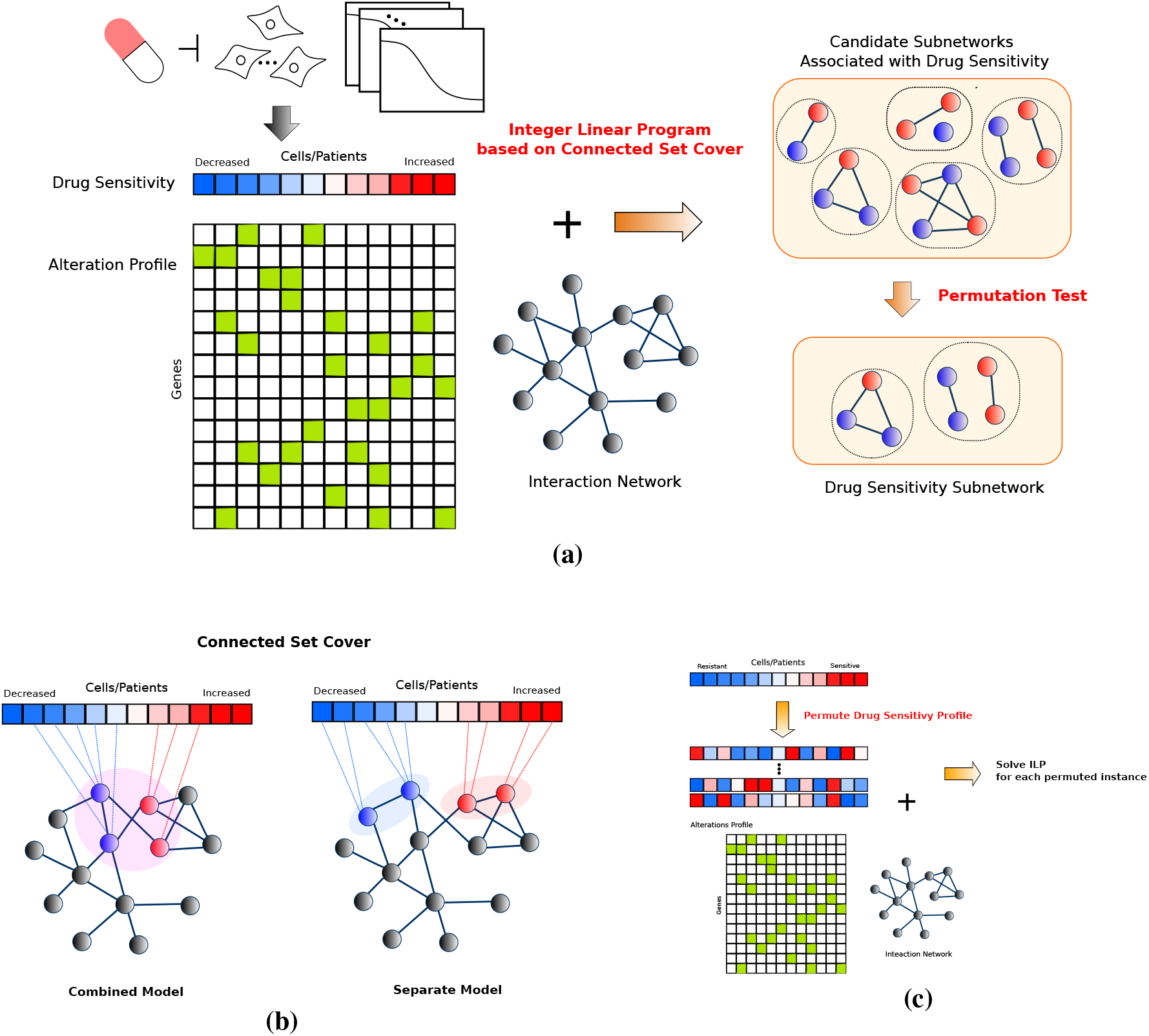
NETPHIX Method. (a) Overview: NETPHIX takes a drug sensitivity profile (continuous phenotype values) and alterations status for the same set of samples as input. In addition, the algorithm utilizes interaction information among genes. Using a connected set cover based ILP algorithm, we first generate a set of candidate modules. The final set of modules are selected based on a permutation test and include only maximal modules among statistically significant optimal solutions. (b) NETPHIX finds a connected set of genes for which corresponding mutations are associated with phenotype values (red colors in the drug response profile indicate increased sensitivity values and blue colors are for decreased sensitivity values). We considered the *combined* model in which all the selected genes are connected regardless of the directions of association and the *separate* model in which two subnetworks are identified for increased and decreased sensitivity module separately (c) The significance of identified modules are assessed using a permutation test in which drug sensitivity profiles are permuted.

#### Connected set cover

For the first step, to obtain candidate subneworks, we design our algorithm based on connected set cover to maximize the total association with drug response (Fig. 1b). Connected set cover approaches have been used successfully for the identification of cancer driving mutations, to overcome the challenges posed by the heterogeneity of cancer mutations and to help uncover relevant genes with rare or medium mutation frequencies [29, 20, 16, 5, 30, 31, 32]. Since we are interested in finding alterations associated with drug response, we seek a connected set of genes that maximizes the total weight where the weights are assigned based on drug sensitivity profile. In addition, to utilize the property of mutual exclusivity, NETPHIX penalizes overlapping mutations when a patient has alterations in more than one selected gene. It has been observed that patient groups harboring different cancer related mutations tend to be mutually exclusive. This property may arise when mutations in two different genes lead to dysregulation of the same cancer pathway and the role of the two genes for cancer progression is redundant. Building on this observation, NETPHIX identifies a connected set of genes *S* such that the sum of phenotypic weights of the patients with alterations in *S* (minus the penalties for overlapping alterations) is maximized. As presented in the results, the penalties for overlapping mutations (i.e., utilizing the property of mutual exclusivity) indeed improves the quality of the solutions compared to the ones we obtained by running NETPHIX without penalties.

There may be genes associated with either directions of drug response - genes whose alterations correlate with increased sensitivity to the drug (decreased cell survival) and genes whose alterations correlate with decreased sensitivity to the drug (increased cell survival). Our algorithm is designed to find a module that includes both types of genes simultaneously. When modeling the connectivity constraints, we considered two different models – the combined model and the separate model (Fig. 1b). In the “combined” model, we identify one connected subnetwork which include all genes associated with either direction, assuming that alterations in different genes belonging to the same functional module can lead to different directions of drug response. In the “separate” model, we seek the solutions with two connected subnetworks, one for increased sensitivity and one for decreased sensitivity separately. This model can capture the case when two different functional modules affect drug response in different ways. As shown later in the results, the combined model finds more associated modules in general than the separate model, although there are a few drugs whose responses are associated more significantly with two separate subnetworks.

#### Selecting final modules

We run multiple ILP instances for different module sizes and connectivity options to obtain candidate modules. Once we obtain the optimal gene module for each parameter combination, we assess the significance of the identified module by performing a permutation test (Fig. 1c). Note that our algorithm is designed to identify the modules associated specifically with a given phenotype (e.g., drug sensitivity to each drug) rather than finding general cancer drivers, and therefore, a permutation test was performed by permuting the drug sensitivity profile so that the significance of the association is assessed in comparison with randomly generated phenotype. Among all significantly associated subnetworks, we obtain the final drug sensitivity modules by selecting maximal modules to remove redundancy. See Section S1.3 for the details of the permutation test and maximal module selection.

### 2.2 Method evaluation on simulated data

We generated a set of simulated instances where we planted phenotype associated modules with varying parameters onto the background of real cancer cell mutation data (Section S1.4). We then compared the performance of NETPHIX and three related methods – LOBICO, UNCOVER and SigMOD. LOBICO is a logic model based algorithm, developed to identify a set of genes whose alterations are related to drug response [23]. UNCOVER [20] was proposed as a method to identify a set of phenotype-associated genes by taking a set cover approach. Both LOBICO and UNCOVER find an optimal solution using an integer linear program but neither algorithms utilizes interaction information. SigMOD is a recently proposed module identification algorithm combining GWAS and network based approach, and was found to outperform other related methods [27]. SigMOD requires individual association scores of genes to a phenotype as an input, for which we used the p-value of association of each gene to a given phenotype by performing t-tests on the coefficients of univariate linear regression.

For the evaluation purposes, we considered simple cases where alterations are associated with only one direction (either increased or decreased). Even though NETPHIX can identify subnetworks with mixed associations simultaneously, UNCOVER considers each direction separately. In addition, the logic models of LOBICO become more complicated and difficult to solve when both sides of associations are present. We planted modules of size 3, 4, and 5 and evaluated the accuracy of the three methods in identifying the planted modules (Fig. 2ab). For all algorithms except SigMOD, we ran the algorithm for different *k*’s (*k* is a parameter for the size of a module searched by the algorithms), while SigMOD automatically adjusted its parameters to find the best module. Also for all ILP based algorithm, we limit the running time up to 24 hours, meaning the algorithms will stop and output the current solution (which may be suboptimal) when the time limit reaches.

**Figure 2:**
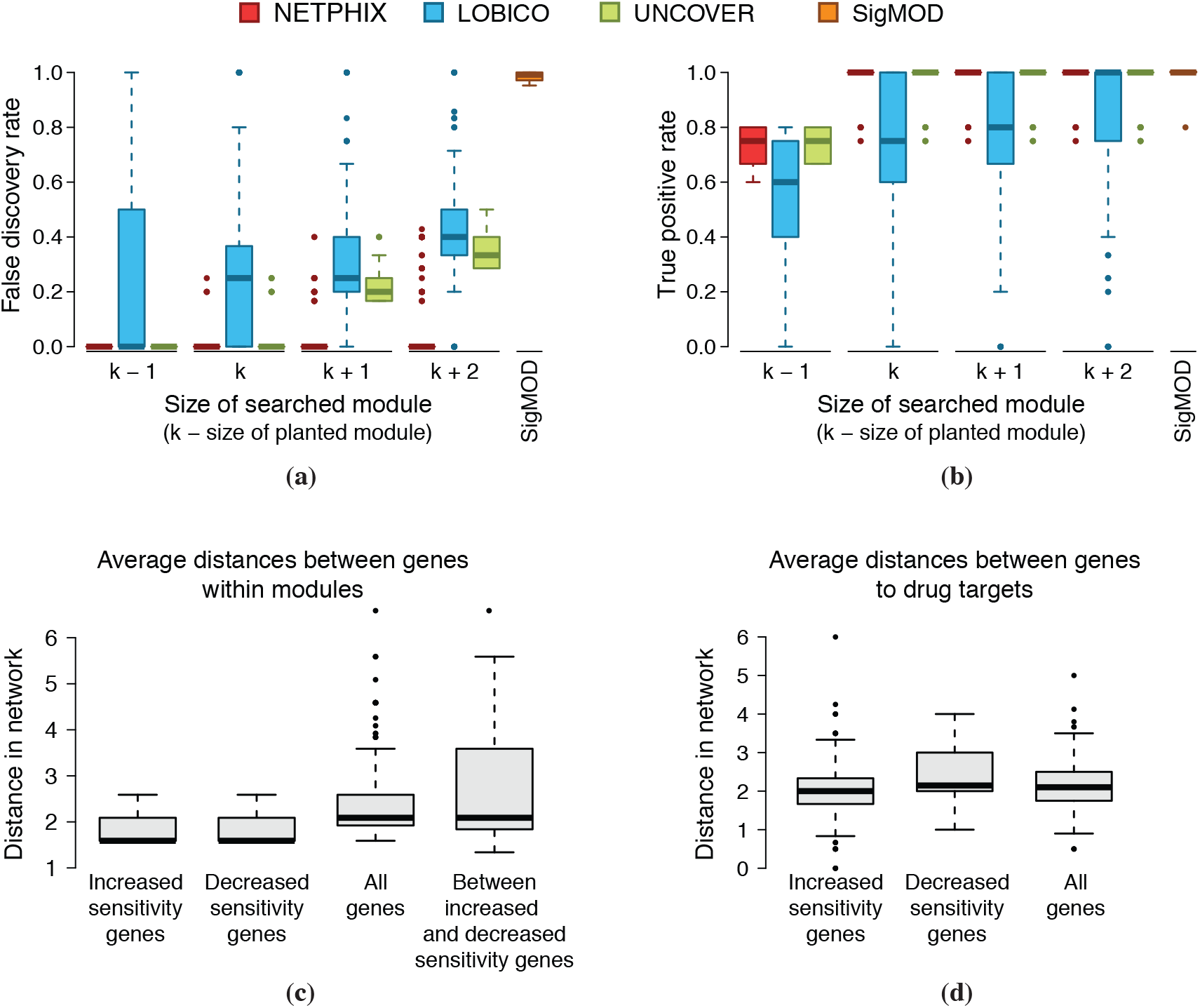
(a-b) Method comparison on simulated data. (a) False discovery rate and (b) True positive rate for the modules identified by NETPHIX (red), LOBICO (blue), UNCOVER (green), and SigMOD (orange). **(c-d) Properties of NETPHIX modules.** (c) Average distances between genes in the selected modules. Distances for genes associated with increased sensitivity (Inc), decreased sensitivity (Dec), all genes in the selected modules (All), and distances between increased sensitivity genes and decreased sensitivity (Between) are shown (d) Average distances between selected genes and the corresponding drug targets. Distances for genes associated with increased sensitivity (Inc), decreased sensitivity (Dec), all genes in the selected modules (All) are shown

As shown in Fig. 2a, only NETPHIX shows very low rate of false positives, i.e., falsely identified genes for all module sizes including when bigger modules than planted are searched. NETPHIX usually does not extend the best module with spurious genes even if we searched for modules bigger than planted while UNCOVER and LOBICO tends to add more genes when increasing *k*. SigMOD identified a large number of false positives along with the planted modules (approx. 100-180 genes) that are not associated with phenotypes. In general, the algorithms uncovered the planted modules in most of instances (Fig. 2b) as long as the size of serached modules are at least as big as the planted module sizes. However, LOBICO solutions miss true positives more often possibly due to the fact that the algorithm returns suboptimal solutions after the time limit reaches. Note that the LOBICO results previously reported have been obtained with pre-selected genes/pathways (consisting of 1,000 elements) while in this simulation we used genome-wide alteration profiles without prefiltering. Both UNCOVER and NETPHIX found optimal solutions well within the time limit (See Fig. S2).

### 2.3 Analysis of drug screening dataset

Using NETPHIX, we analyzed a large scale drug screening dataset (GDSC) for which genomic alteration profiles for hundreds of cell lines and drug sensitivity data for 265 drugs are available (Section S1.4). Application of NETPHIX to the dataset resulted in identifying a total of 641 modules for 203 drugs (for the remaining drugs no modules with significant association were identified, Table S1). Since there can be multiple functional modules affecting drug efficacy, our method allows to identify multiple associated modules for a specific drug. Out of 641 identified modules, 335 modules are one connected modules based on the combined model (for 183 drugs) and 306 modules consist of two connected components based on the separate model (for 171 drugs). See Section S1 for detailed description on how the final modules are selected.

#### NETPHIX modules are functionally related and close to the drug target

NETPHIX is designed to choose modules in which genes are highly connected. As a result, we observed that the genes in the selected modules are close in terms of distance in the network (the mean of avg. distances = 1.72, Fig. 2c), suggesting that the genes are closely related functionally. In addition, the genes with the same direction of association (decreased or increased) tend to be closer to each other than the genes associated in opposite direction although the two groups of genes are still close in the network (the mean of avg. distances between the two groups = 1.99).

To examine the relationship between the sensitivity modules and the targets for the corresponding drugs, we also computed the distances between the drug targets and the selected genes for each drug (Fig. 2d). We found that the genes in drug sensitivity modules are located near drug targets in the network (the mean of avg. distances = 2.13). Interestingly, we observed that the genes associated with increased sensitivity are closer to drug targets than the genes associated with decreased sensitivity (*p <* 10^*−*13^, *t*-test), indicating that having perturbations in genes closer to drug targets could potentially improve the efficacy of the drugs.

#### NETPHIX identified biomarkers for drugs

Many of the modules identified by NETPHIX provide interesting insights related to drug action. In particular, we analyzed the response to drugs targeting the RAS/MAPK pathway (Fig. 3e). This pathway regulates growth, proliferation and apoptosis and is often dysregulated in various cancers. Among the most common mutations of this pathway are mutations of BRAF/KRAS/NRAS. Interestingly, NETPHIX identified modules including those genes (mostly BRAF, KRAS and sometimes NRAS) as associated with increased sensitivity to all MEK inhibitors (Selumetinib, Trametinib, CI-1040, PD0325901, and Refametinib) and an ERK inhibitor (VX-11e). All these six drugs act by blocking MEK1/MEK2 or ERK genes that are immediately downstream of BRAF/KRAS/NRAS and the increased sensitivity attributed to the alterations in this subnetwork is consistent with the action of these drugs. Modules associated with decreased sensitivity to the drugs are more diverse but NETPHIX frequently selected the module of genes ERBB2 (amplification), MYC and RB1 (mutations) or the module with TP53 mutations. All the genes in the modules are related to the MAPK/ERK signaling pathway. The mutation status of BRAF and KRAS, the core members of the pathway, were previously identified as predictors of MEK inhibitors although KRAS mutations can affect drug responses differently depending on the mutation types [33, 34, 35]. ERBB2 is a receptor protein that signals through this pathway, while MYC, RB1 and TP53 are downstream of the MAPK/ERK signaling pathway. RB1 was found to be associated to the resistance to MEK inhibitors [36] and MYC degradation by inhibition of MEK leads to an increase in both ERBB2 and ERBB3 mRNA expression, causing intrinsic drug resistance [35]. TP53 mutations are associated with multiple drug resistance [37, 38]. These findings indicate that the alterations in different components of the same pathway can contribute to drug sensitivity in different ways.

**Figure 3:**
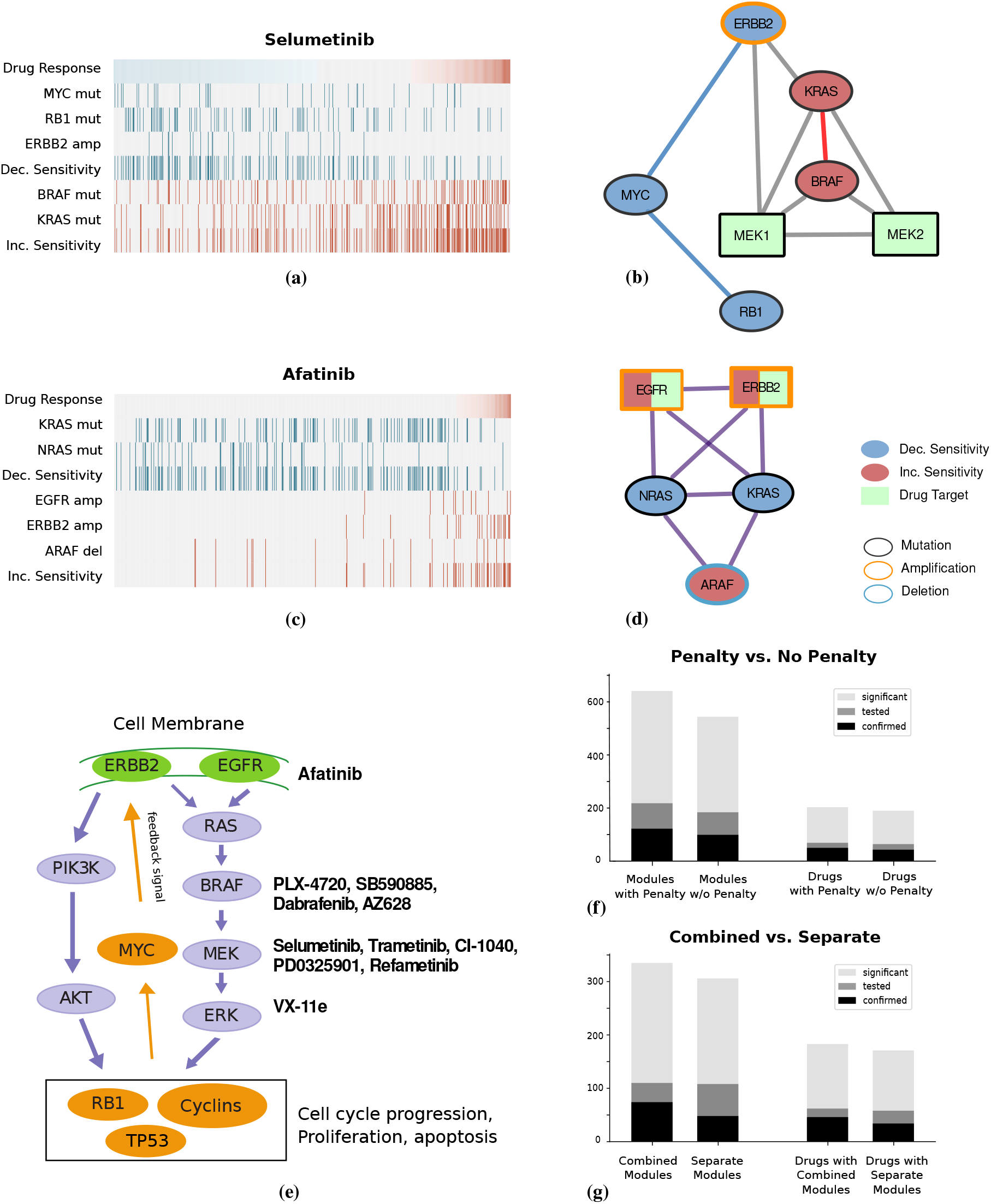
(a-d) Sensitivity networks identified by NETPHIX. Alternation profile and connectivity between the drug target and the genes for Selumetinib (a-b) and Afatinib (c-d). In the alteration profile, the panel shows the values of the phenotype (i.e., drug response, top row) for all samples (columns), with blue being decreased sensitivity values and red being increased sensitivity values. For each gene, alteration status in each sample are shown in red/blue (with the summary covers for decreased and increased sensitivity separately), while samples not altered are shown in grey. The module for Selumetinib is identified based on the separate connectivity model and the module for Afatinib is selected based on the combined connectivity model. **(e) Schematic diagram of MAPK/ERK and AKT signaling pathways** with drugs and their drug targets annotated. **(f-g) Comparison between different options.** (f) Comparison between runs with and without penalty. The number of significant modules/drugs (Sig), the number of tested modules/drugs with CTRP (Tested) and the number of confirmed modules/drugs with CTRP (Validated) are shown. Drugs are counted when there is at least one associated modules that are significant, tested or validated, respectively. (g) Comparison between the combined and separate connectivity models.

In contrast to the response to MEK1/2 and ERK2 inhibitors, the drugs directly targeting BRAF are associated with more heterogeneous modules. While all BRAF inhibitors (except HG6-64-1) commonly exhibit increased sensitivity in BRAF mutant cell lines, KRAS mutations help the action of Type II BRAF inhibitors (AZ628) but develop resistance to Type I inhibitors such as Dabrafenib, PLX-4720, and SB590885, which is consistent with the previous findings [39]. This suggests that patient specific mutational profiles can provide important clues in predicting drug response.

#### NETPHIX modules suggest candidates for drug combination therapy

We hypothesize that pairs of drugs can potentially be candidates for combination therapy if they are associated with similar modules but the genes in the modules are associated with drug responses in opposite directions. By analyzing the identified modules for pairs of drugs with such property, we identified 237 drug pairs (Table S3). Although the systematic validation could not be performed due to the lack of validation dataset, we found the evidences in literature for the efficacy of several drug combinations. For example, Afatinib, a pan-ErbB inhibitor, have associated modules of KRAS, NRAS (mutations) for decreased sensitivity and EGFR, ERBB2 (amplification) and ARAF (deletion) for increased sensitivity. This suggests that it might be beneficial to use Afatinib in combination with MEK 1/2 and ERK2 targeting drugs discussed above (Fig. 3a-d). Indeed, studies showed that Afatinib and Selumetinib work synergistically [35] and clinical trials for combination therapy are currently underway [40]. In addition, the efficacy of PD0325901 and Afatinib combination is also reported [41]. Both Selumetinib and PD0325901 have associated modules similar to Afatinib but in opposite direction.

Another example is the combination of Lapatinib and Vorinostat (Fig. S1cd). Lapatinib is a drug that inhibits EGFR/ERBB2 and Vorinostat is a histone deacetylase (HDAC) inhibitor. Vorinostat has been shown to improve how well Lapatinib kills cancer cells in clinical trials [42] and Lapatinib in general enhances the antitumor activity of the histone deacetylase inhibitor synergistically [43].

### 2.4 Validation of identified sensitivity modules with independent datasets

To examine if the alterations in the modules identified by NETPHIX indeed lead to different responses for the corresponding drugs, we performed the validation of the identified subnetworks with CTRP (Cancer Therapeutics Response Portal), an independent drug screening data. Among 203 drugs for which NETPHIX identified at least one module, 69 drugs have the drug response profiles reported in CTRP dataset, and many of the drugs have multiple associated modules, resulting in 218 modules available for validation. We divided the cell lines in the dataset depending on the alteration status in the identified genes for each drug and tested if the cell survival rates differ between different groups (Table S2). We found that 50 out of 69 drugs (72%) have at least one module having a statistically significant difference (*p <* 0.05 (ANOVA), FDR *<* 9% (BH)), and 122 modules in total (out of 218, 56 %) were confirmed in the validation.

### 2.5 Impact of NETPHIX design choices on the results

While the benefits of using gene interactions for the identification of phenotype associated genes are generally accepted, the usage of mutual exclusivity in this context has not been investigated before. Similarly, the question whether it is better to model the decreased and increased sensitivity genes in one combined module or two separate modules need to be also examined. Therefore, we use the same validation set to compare the performance of different options of our algorithm.

#### Mutual exclusivity helps identify drug sensitivity modules

Our objective function in ILP includes penalties for overlapping mutations, which lets NETPHIX preferentially select genes whose alterations are mutually exclusive. To investigate the effects of using penalties on the performance, we ran NETPHIX *without penalties* in the objective function and compared with our results obtained when penalties are used. As shown in Fig. 3f, NETPHIX finds a smaller number of significant modules when penalties are not included (544 modules compared to 621 modules in the original solution). The number of drugs with at least one associated modules is also smaller (190 drugs without penalty vs. 203 drugs with penalty). We further compared the effects of penalties in terms of the number of drugs/modules confirmed in CTRP dataset. Without penalty, only 99 modules out of 184 tested modules (54%) were confirmed whereas 122 modules out of 218 tested modules (56%) were confirmed when penalty were used. In addition, 43 drugs had at least one confirmed modules (67% of tested drugs) without penalty compared with 50 drugs with penalty (72% of tested drugs). Overall the results show that mutual exclusivity helps find more modules that are statistically significant, and the identified modules has a higher percentage of true positives as demonstrated in the validation using the independent dataset.

#### Combined vs. separate connectivity model

Next, we compared the performance of two connectivity models – the combined model where all selected genes are connected and the separate model in which two modules identified for increased and decreased sensitivity separately (Fig. 3g). Slightly more modules were identified with the combined model than with the separate model (335 vs. 306 modules), and more drugs have at least one significant module in the combined model than the separate model (183 vs. 171 drugs). The combined model also has a higher percentage of confirmed modules/drugs. Among 110 and 108 modules tested with CTRP dataset for the combined and separate model respectively, 74 (67%) and 48 modules (44%) were confirmed. In terms of the number of drugs with at least one confirmed modules, the combined model has 46 out of 62 drugs (74%) confirmed whereas the separate model has 34 drugs confirmed out of 58 (59%).

Among the drugs with only the combined model being confirmed in CTRP are IGF-1R inhibitors, BMS-536924 and BMS-754807 (Another IGF-1R inhibitors, Linsitinib does not have the screening data available in CTRP). The modules commonly have KRAS mutations associated with increased sensitivity and PTEN (mutations and deletions) with decreased sensitivity (Fig. S1a). Both genes have been previously shown as biomarkers for IGF-1R inhibitors [44, 45, 46, 47] although there are conflicting reports depending on cancer types and molecular status of other genes [48, 45]. The module that NETPHIX identified for BMS-754807 was confirmed in the CTRP dataset (Fig. S1a), showing a significant different in the cell survival rates of the two groups (*p <* 0.00035, ANOVA).

However, some drugs such as Cytarabine have modules in the combined and separate model, both of which are confirmed in CTRP dataset. In particular, the module associated with Cytarabine in the separate model (Fig. S1b) includes UGT2B17 amplification and CYP2E1 deletion associated with decreased sensitivity. Both enzymes are hypothesized to be an important player in the metabolism of common drugs [49, 50].

## 3 Discussion

We developed a new computational method, NETPHIX (NETwork-to-PHenotpe assocIation with eXclusivity), for the identification of mutated subnetworks that are associated with a continuous phenotype. Using simulations and analyzing a large scale drug screening dataset, we showed that NETPHIX can uncover the subnetworks associated with response to cancer drugs with high precision. We found many statistically significant and biologically relevant modules associated with drug response, including MAPK/ERK signaling related modules associated with opposite response to drugs targeting RAF, MEK and ERK genes. The genetic alteration status in many of identified modules indeed make differences in cell survival rates, as validated with an independent dataset. Overall, the modules identified by NETPHIX are in good correspondence with the action of the respective drugs, suggesting that NETPHIX can correctly identify relevant modules and the modules can thus be used to predict potential patient-specific drug combinations and to provide guidance to personalized treatment.

We demonstrate that the preferential selection of mutually exclusive genes was important for a better performance of the method. Interestingly, although one might assume that genes affecting drug resistance are not necessarily functionally related to the genes increasing drug sensitivity, we found that the combined connectivity model outperforms the separate connectivity model, indicating that the two groups of genes in fact might be related.

The applicability of NETPHIX can go far beyond the drug response discussed in this paper, to any continuous cancer phenotypes. We expect that NETPHIX will find broad applications in many types of network-to-phenotype association studies.

## Acknowledgements

This research was supported in part by the Intramural Research Programs of the National Library of Medicine at National Institutes of Health, USA. FV was supported, in part, by the University of Padova grants “SID2017” and “STARS: Algorithms for Inferential Data Mining”. We would like to thank Jan Hoinka for helpful discussions.

## Supplementary Materials

### S1 Methods

#### S1.1 Formal definition of the computational problem

We are given a graph *G* = (*V, E*), with vertices *V* = {1,…, *n*} representing genes and edges *E* representing interactions among genes. Let *P* denote the set of *m* patients (or cell lines). For each sample *j* ∈ *P*, we are also given a phenotype profile value *w*_*j*_ ∈ ℝ which quantitatively measures a phenotype (e.g., drug response in our study). Let *P*_*i*_ ⊆ *P* be the set of patients in which gene *i* ∈ *V* is altered. We say that a patient *j* ∈ *P* is *covered* by gene *i* ∈ *V* if *j* ∈ *P*_*i*_ i.e. if gene *i* is altered in sample *j*. We say that a sample *j* ∈ *P* is *covered* by a subset of genes (or vertices) *S* ⊆ *V*, if there exists at least one vertex *v* in *S* such that *j* ∈ *P*_*v*_.

For simplicity of description, we start with the formulation in the case where the association is in one direction, for example, with increased drug sensitivity. Later we will show how to extend the problem to accommodate the case where mixed associations are allowed in the same module. Our goal is to identify a connected subgraph *S* of *G* of at most *k* vertices such that the sum of the weights of the samples covered by *S* is maximized. The weights are computed based on drug sensitivity. Since we are interested in functionally complementary mutations, we also penalize coverage overlap when a sample is covered more than once by *S* by assigning a penalty *p*_*j*_ for each of the additional times sample *j* is covered by *S*. Let *c*_*S*_(*j*) be the number of times element *j* ∈ *P* is covered by *S*. For a set *S* of genes, we define its weight *W* (*S*) as:

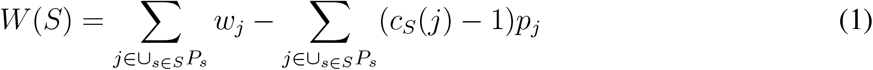

Thus, we define the optimization problem for one-side association as follows: Given a graph *G* defined on a set of *n* vertices *V*, a set *P*, a family of subsets *P* = {*P*_1_,…, *P*_*n*_} where for each *i*, *P*_*i*_ ⊆ *P* is associated with *i* ∈ *V*, weights *w*_*j*_ and penalties *p*_*j*_ ≥ 0 for each sample *j* ∈ *P*, find the subset *S* ⊆ *V* of ≤ *k* connected vertices maximizing *W* (*S*).

Since genetic alterations may affect the increase or decrease of drug sensitivity, we extend the problem to identify genes with associations in both directions in one module. Considering genes with increased and decreased sensitivity simultaneously can pick up stronger signals of associations and allow to take into account the interactions between alterations affecting drug responses in different ways. Let *I* include the genes associated with increased sensitivity overall (i.e., genes *i* with positive total weights, 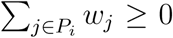) and *D* is the set of genes associated with decreased sensitivity overall (i.e., genes *i* with negative total weights, 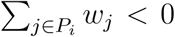). Our objective function is then defined as follows:

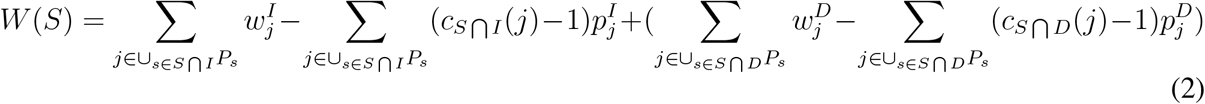

where we define 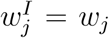 and 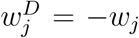. We considered two versions of connectivity constraints among the associated genes as illustrated in Fig. 1b. In the first model, we insisted that all selected genes should be connected whether they are associated with increased or decreased sensitivity. In the second model, we ensured the connectivity of genes with the same direction of association, resulting in two connected components in a solution (one for increased and the other for decreased sensitivity).

Although the problem is NP-hard (by a reduction to set cover) even for the simple one-sided case without network constraints, we formulated it as an integer linear program as described in the next subsection, and solved it to optimality using CPLEX, which can be run in a reasonable amount of time (See Fig. S2 for running times for different *k*’s).

#### S1.2 ILP formulation

Let *x*_*i*_ be a binary variable (denoted with *x*_*i*_ ∈ 𝔹) equal to 1 if gene *i* ∈ *V* is selected and *x*_*i*_ = 0 otherwise. Let 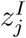(resp., 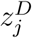) be a binary variable equal to 1 if sample *j* is covered by a gene *i* ∈ *I* (resp., *i* ∈ *D*) and 0 otherwise. Let 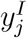(*resp*., 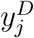) denote the number of genes in *I* (resp., *D*) cover sample *j* in the solution. Finally, let *w*_*j*_ be the weight of sample *j* and *p*_*j*_ be the penalty for sample *j*. When sample *j* is covered by a gene in *I*, the weight and penalty remain the same 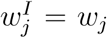. When *j* is covered by a gene in *D*, 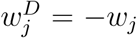. Our ILP formulation for the combined model is defined as follows:

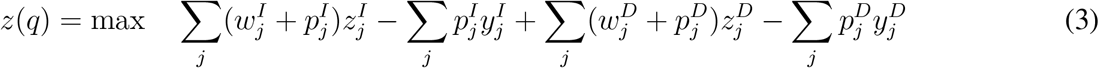

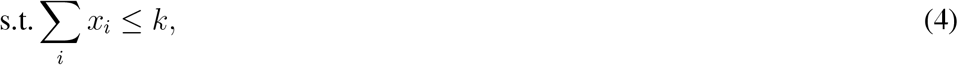

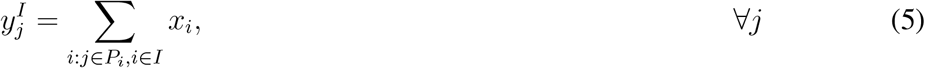

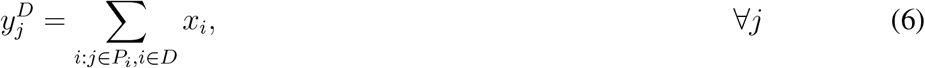

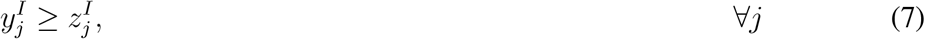

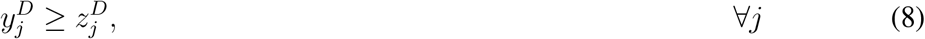

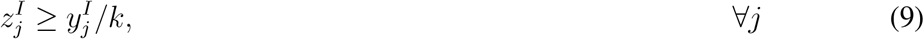

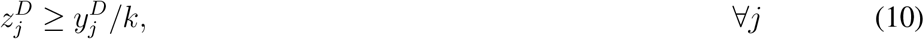

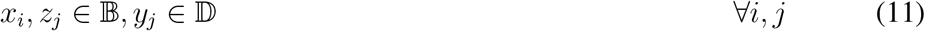

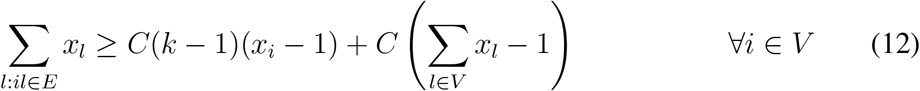

Constraint (4) impose that the total number of sets (i.e., selected genes) in the solution is at most *k*. Constraints (5) and (6) define how many times each sample has been covered by genes in *I* and *D*, respectively. Constraints (7) (resp., Constraints (8)) ensure that for each sample *j* ∈ *P*, if *j* is covered by increased (resp., decreased) sensitivity genes in the current solution then the number of times *j* is covered by *I* (resp., *D*) in the solution is at least 1. Constraints (9) (resp., Constraints (10)) impose that for each element (sample) *j* ∈ *P*, if *j* is covered by at least one increased (resp., decreased) sensitivity gene in the current solution then *j* is covered by *I* (resp., *D*).

Constraints (12) were used to ensure the high connectivity of a selected module (the combined connectivity model). Specifically, the constraints enforce that each selected gene is connected with at least *C* fraction of genes in the selected module (other than the gene itself). Note that if *C* ≥ 0.5, the module is a connected subgraph since for any two non-adjacent vertices, they must have a common neighbor (*C* = 0.5 is used in our analysis). In our study, we used a functional interaction network (from STRING database), which is relatively dense. For sparse networks where highly connected components are rare, we may use an alternative approach based on a branch-and-cut algorithm to ensure the connectivity [51, 52, 53].

Note that Constraints (12) forces the connectivity among all selected genes regardless of the directions of association. For the separate connectivity model, we identify candidate modules so that the connectivity is only enforced among the genes in *I* and *D*, separately. In this case, we replace the connectivity constraints given in (12) with the following constraints.

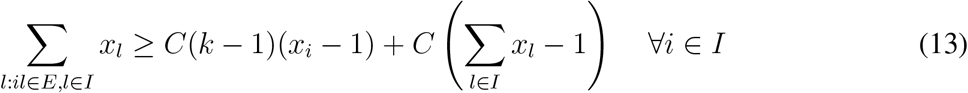

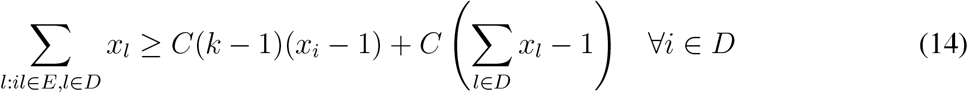

#### S1.3 Selecting final modules

By computing the optimal solutions of ILP instances with different sizes *k* (*k* = 1 to 5) and two connectivity options (the combined and separate model), we first obtain a pool of candidate modules. For each candidate module, we run a permutation test to assess the statistical significance of association and select maximal modules among significantly associated ones. Note that we allow to choose multiple modules associated with a drug in the final solution because it is possible that multiple functional components are associated with drug response.

##### Permutation test

For each candidate module, we assess the statistical significance of the association between their alteration profile and drug response by a phenotype permutation test. In the phenotype permutation, the dependencies among alterations in genes are maintained, while the association between alterations and the phenotype is removed. Specifically, a permuted dataset under the null distribution is obtained as follows: the graph *G* = (*V, E*) and the sets *P*_*i*_, *i* ∈ *V* are the same as observed in the data; the values of the phenotype are randomly permuted across the samples (Fig. 1c). To estimate the *p*-value for the solutions obtained by ILP, we used the following standard procedure: 1) we run an algorithm on the real data *𝒟*, obtaining a solution with objective function *o_𝒟_*; 2) we generate *N* permuted datasets as described above; 3) we run the same algorithm on each permuted dataset; 4) the *p*-value is then given by (*e* + 1)*/*(*N* + 1), where *e* is the number of permuted datasets in which our algorithm found a solution with objective function ≥ *o*_*𝒟*_. We used *N* = 100 permutations in our analysis and considered the modules with *p <* 0.01 (FDR *<* 3%, BH) as significantly associated modules.

##### Selecting maximal modules

Among all significantly associated modules obtained based on the permutation test, we remove redundant modules by selecting only maximal modules. In other words, let *M*_1_, *M*_2_, …, *M*_*t*_ be the set of significantly associated modules for a drug. For any two modules *M*_*i*_ and *M*_*j*_ such that *M*_*i*_ ⊂ *M*_*j*_ then we only include *M*_*j*_ in the final solution for the drug.

#### S1.4 Datasets and Method Details

##### Drug sensitivity dataset

The Genomics of Drug Sensitivity in Cancer Project (https://www.cancerrxgene.org/) consists of drug sensitivity data generated from high-throughput screening using fluorescence-based cell viability assays following 72 hours of drug treatment. In particular, we considered the area under the curve for each experiment as a phenotype. These scores are provided in the file `portal-GDSC_AUC-201806-21.txt` available through the DepMap data portal (https://depmap.org) for 265 compounds and 743 cell lines, with 736 having alteration data available through the DepMap portal. For the DepMap experiments [54, 55], we used the alteration provided at https://depmap.org/portal/download/all/. We downloaded the data on July 6^*th*^ 2018. In particular we used mutation data from the file `portal-mutation-2018-06-21.csv` that includes binary entries for 18,652 gene-level mutations. Additionally, we considered 22,746 amplifications and 22,746 deletions computed from the gene copy number data in `portal-copy_number_relative-2018-06-21.csv`, with an amplification defined by a copy number above 2 and a deletion defined by a copy number below −1.

##### Preprocessing drug sensitivity data

For every drug response profile, we excluded samples with missing values for that pheno-type, which results in a different number of samples for each phenotype. The number of samples varied between 240 and 705. To generate drug sensitivity values for the patients, we took the negatives of cell viability (i.e., increased cell survival indicates decreased sensitivity to the drug and vice versa) and then normalized the phenotype values before running the algorithm, by using standard z-scores (subtracting the average value 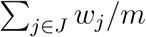 from each weight *w*_*j*_ and dividing the result by the standard deviation of the (original) *w*_*j*_’s), in order to have both positive and negative phenotype values. We excluded genes with low (present in less than 1% samples from our analyses. As penalty for increased sensitivity 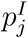, we use the average of the positive phenotype values if the original value of the element was positive (*w*_*j*_ > 0) and assign a penalty equal to its absolute value otherwise. The penalty for decreased sensitivity 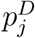 is computed in the opposite way. The negative of the average of the negative phenotype values is used if the original value of the element was negative (*w*_*j*_ < 0) and assign a penalty equal to its absolute value otherwise

##### Interaction network and computing distances in the network

For functional interactions among genes, we used the data downloaded from STRING database version 10.0 [56]. We only included the edges with high confidence scores (≥ 900 out of 1000) as an input to NETPHIX. The resulting interaction network includes 9,215 nodes and 160,249 edges.

For the average distances within modules, we computed the pairwise shortest distances within modules and take the average distances. To compute the distances from selected modules to drug targets, we used the drug target information given in Table S1. For each drug, we only used the drug targets present in the functional network that are reachable from the selected modules and computed the average distance for all pairs of genes.

##### Running simulated experiments

For the background of simulation data, we use the same gene alteration table and interactions from drug sensitivity dataset described previously in this section. The phenotype values for individual samples are randomly drawn from normal distribution *N* (0, 1). We then planted randomly generated phenotypes and associated modules to the background as follows.

Phenotypes: *α* fraction of patients *P* (*α*) (*α* = 0.1, 0.2, and 0.3) were randomly selected and assigned phenotype values drawn randomly from *N* (*z*, 0.5) where *z* is a z-score corresponding to a cumulative p-value *p* (*p* = 0.005, 0.1, 0.99, and 0.995).

Associated gene modules: we randomly selected a gene set *S*(*k*) of size *k* (*k* = 3, 4, and 5) and added random alterations in *S*(*k*) for patients *P* (*α*) so that each patient in *P* (*α*) has an alteration in exactly one gene in *S*(*k*). Therefore, the added alterations among the patients *P* (*α*) are mutually exclusive although there may be overlapping mutations due to the background alterations. We also added random edges among the genes *S*(*k*) so that they satisfy the density constraints (*C* = 0.5)

We generated 10 random instances for each combination of parameters (*k*, *α*, *z*) and ran the module identification algorithms.

For LOBICO [22], we used its R implementation [57] with the default parameter settings, except the logic function parameters (*K* and *M*) and the maximum running time. The OR logic model with *K* = *k* and *M* = 1 was used for increased sensitivity modules and the AND logic module with *K* = 1 and *M* = *k* for decreased sensitivity modules, where *k* is the size of the searched module. We limited the running time of LOBICO to be 24h and reported the best current solution (which may be suboptimal) when the program stops.

##### Validation of identified modules with CTRP dataset

For validation of NETPHIX modules, we utilized an independent drug response dataset from the Cancer Therapeutics Response Portal (CTRP) [58]. The drug screening results were downloaded from https://portals.broadinstitute.org/ctrp/ (Version 2). The area under the curve (AUC) values were used for drug response phenotypes. For the alteration profiles for the cell lines, we used `CCLE_MUT_CNA_AMP_DEL_binary_Revealer.gct` downloaded from https://portals.broadinstitute.org/ccle/data (08/21/2017).

We found the drug response profiles for 76 drugs in both CTRP and GDSC datasets, among which 69 drugs have at least one drug sensitivity module identified by NETPHIX. 821 cell lines having both drug response and gene alteration profiles were used for validation. To test if the alteration status of selected genes are associated with different drug response, we divided the cell lines into three groups; The cell lines (*C*_*I*_) with alterations in increased sensitivity genes but no alterations in decreased sensitivity genes, the cell lines (*C*_*D*_) with alterations in decreased sensitivity genes but no alterations in increased sensitivity genes, and the cell lines (*C*_*N*_) with no mutations in the identified genes. We then performed ANOVA test for the cell survival rates for the three groups (*C_I_, C_D_*, and *C*_*N*_).

**Figure S1:**
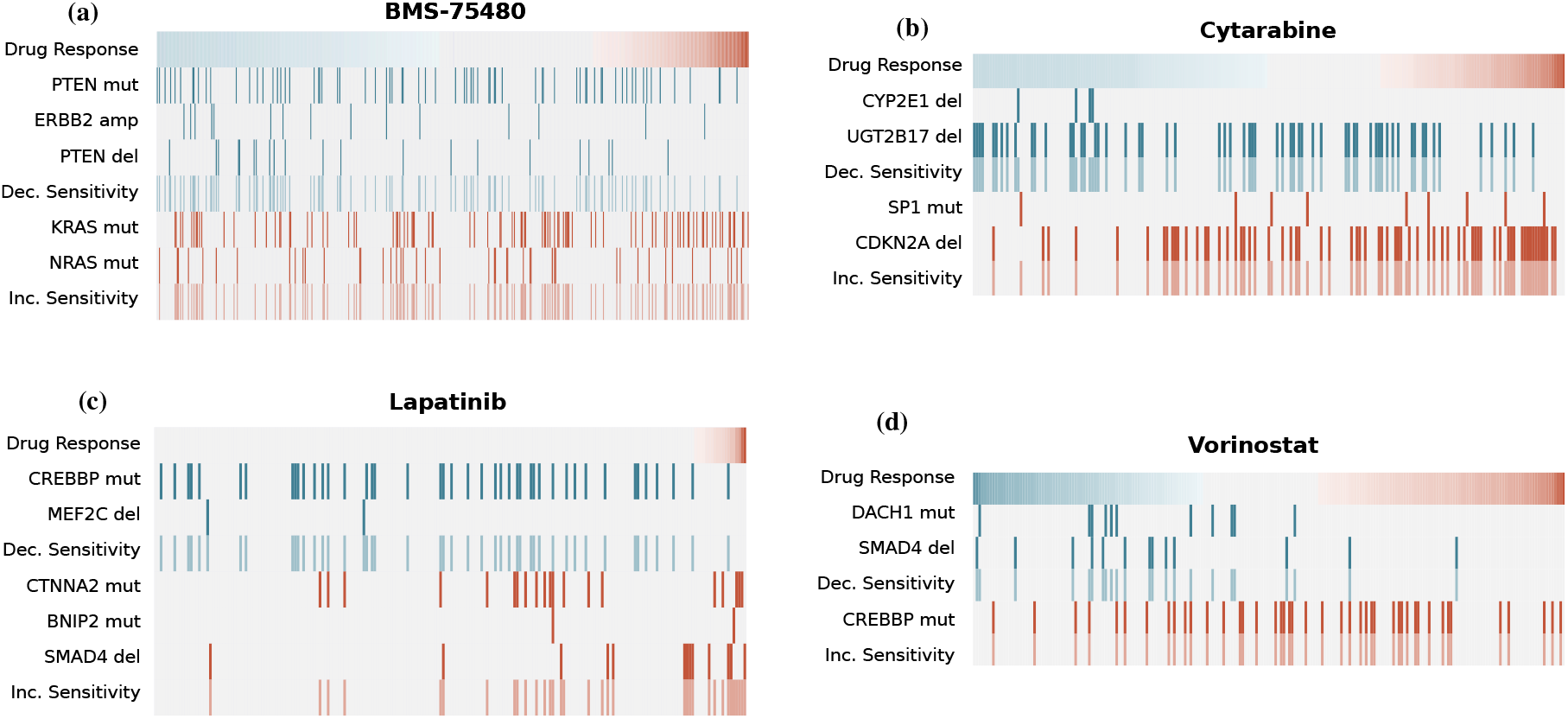
Modules identified by NETPHIX. (a) Sensitivity module for Cytarabine identified based on the combined connectivity model. (b) Sensitivity module for JQ1 identified based on the separate connectivity model. (c-d) Sensitivity module for Lapatinib (c) and Vorinostat (d). The two modules associated with the drugs are similar but they are associated with opposite directions. The efficacy of combination therapy with Lapatinib and Vorinostat is confirmed in clinical trials.

**Figure S2:**
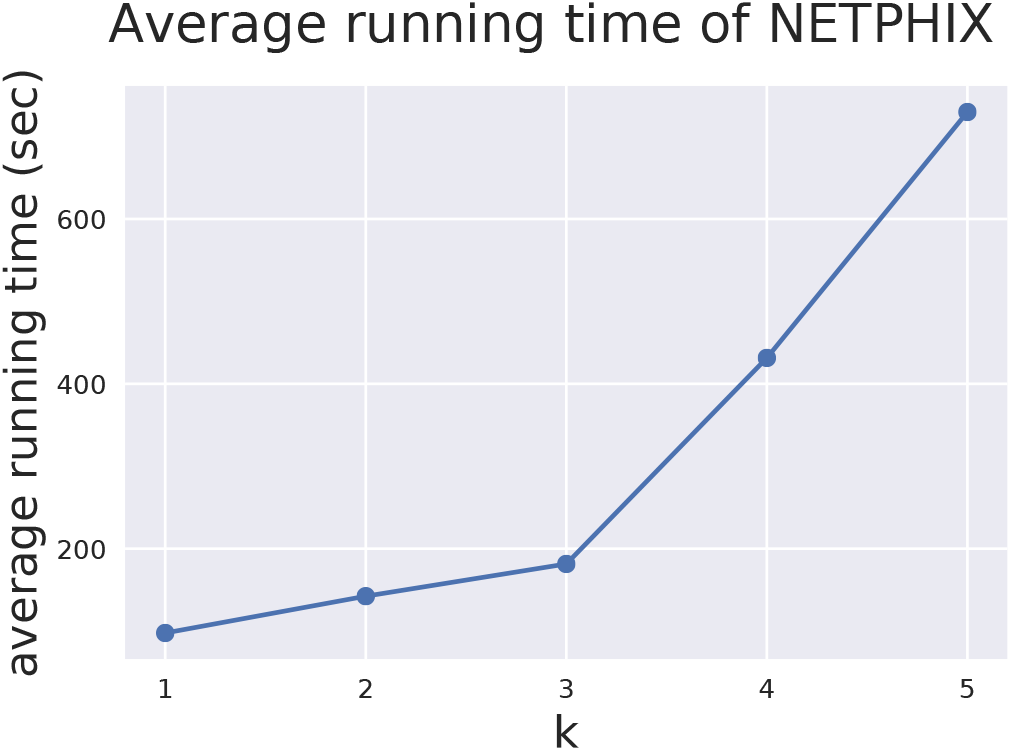
The average running times of NETPHIX over different *k*’s

